# *Borrelia burgdorferi* 297 *bmpA* encode the mRNA that contains ORF for a leader peptide that regulates *bmpA* gene expression

**DOI:** 10.1101/542589

**Authors:** L.P Dubytska

## Abstract

The Bmp proteins are highly conserved proteins with no well established functions in *B. burgdorferi* sensu lato and are immunogenic. It was reported that four genes from this cluster *bmpD-bmpC-bmpA-bmpB* are expressed *in vitro* as monocistronic and polycistronic messages.

Evidence is presented in this report that *bmpA* mRNA contains two ribosome binding sites (SD) separated by 90 bases pairs. The SD_1_ precedes a small 32 amino acid ORF - leader peptide (BmpAL). The SD_2_ is the RBS for 342 amino acids BmpA. The *bmpA*_*L*_ and *bmpA* ORFs in *B.burgdorferi* 297 overlap by eight base pairs suggesting that two proteins can be co-regulated. First five codons in the leader peptide and “-<u>GGG-“</u> in SD_2_ are rarely used in *Borrelia*, suggesting that they can regulate BmpA_L_ and BmpA expression. Deletion of SD_1_ in the leader sequence, or introducing a stop codon immediately before SD_2_ leads to increased BmpA::GFP expression in *B.burgdorferi* 297 that contains *bmpA::gfp* translational fusion on the plasmid. In *B. garinii* G25 and *B. afzelii* IP3 the leader sequence is in frame with *bmpA*, and a result, in *B. afzelii* IP3 BmpA are expressed as the higher molecular weight protein compared to BmpA of *B. burgdorferi* 297 and *B. afzelii* DK7.

## Introduction

*Borrelia burgdorferi*, the spirochetal bacterium that causes the tick-borne infection called Lyme disease [1, 2]. *B. burgdorferi* genome contains approximately 1000 chromosomal and 400 plasmid genes [3] but only a few homologs to regulatory genes, sigma factors and one *rho* terminator factor [3]. In addition, *Borrelia* has genes and gene families that do not share homology with genes of other bacteria [3] suggesting that *B. burgdorferi* may have different mechanisms to control gene expression.

Evolutionary selected systems of virulence gene regulation allow coordinated gene expression that is based on the temporal and special requirements of host niches. Global regulation of virulence genes is a common strategy of bacterial pathogens to overcome the complexity of innate host defenses [4–11]. In addition to a global regulatory system, prokaryotes can employ non-global mechanisms of virulence gene regulation. They include expression of non-coding RNAs [12, 13], effects on mRNA secondary structure that forms terminator/antiterminator structure [14–16] and affects mRNA stability [17] as well as the differential efficiency of ribosomal binding [18, 19].

The *bmp* gene cluster of *B. burgdorferi* is located in the chromosome and encodes lipoproteins with high amino acid homology, that are expressed *in vivo* and are immunogenic [20–22]. In humans and animals antibodies against one of the members of this family, BmpA (formerly p39), appear early during infection [21]. *B. burgdorferi* with *bmpA* or *bmpB* deletions is unable to persist in mouse joint tissues [23]. The BmpA can also stimulate the production of inflammatory cytokines in human and murine lymphocytes, indicating an important role of BmpA in the maintenance of mammalian infection [23].

According to Dobricova et al. [24], four *bmp* genes are expressed *in vitro* and constitute two transcriptional units with a complex pattern of transcription, including alternative monocistronic and polycistronic messages. One unit contains *bmpD*, and the second unit includes *bmpC, bmpA* and *bmpB*. Moreover, promoters were identified for *bmpD*, *bmpC* and *bmpA*, but not for *bmpB*. The *bmpC* is always expressed as a polycistronic message with *bmpA*, and *bmpA* can transcribe as individual mRNA and as bicistronic *bmpA-bmpB*. According to Ramamoorthy et al. [25] expression from the *bmpA-bmpB* operon results in three distinct transcripts *bmpA, bmpA-bmpB* and *bmpA* truncated *bmpB*. In addition, the conservation of *bpm* genes within the *B. burgdurferi sensu lato* complex and the presence of orthologs in *Treponema pallidium* and numerous other bacteria suggest that these proteins can play an essential physiological role.

Unusual genetical structure Bmp genes and pattern of their expression may indicate specific regulatory mechanisms that are involved in the expression of these genes. To uncover some of the questions about BmpA expression and regulation, we investigate *bmpA* transcript and role of the leader sequence (*bmpA*_*L*_) on BmpA expression.

## Materials and methods

### Bacterial strains and medium

*E. coli* DH5α (New England BioLabs, Beverly, MA) and *E. coli* TOP10 were grown in Luria-Bertani (LB) broth or plates (Gibco-BRL, Gaithersburg, MD). The *B. burgdorferi* 279 [26] was grown in BSK-H medium (Sigma, St. Louis, MO.) with 6% rabbit serum (Sigma, St. Louis, MO). Appropriate antibiotics were added when specified.

**DNA manipulations** were performed by standard methods [27]. Restriction enzymes were obtained from New England BioLabs, Beverly, MA. Total DNA was purified from bacterial cultures using High Pure PCR Template Preparation kit (Roche, Mannheim, Germany), DNA fragment and PCR product purification was done using QIAquick Gel Extraction kit (Qiagen, Valencia, California.); all methods were performed according to the manufacturers’ instructions. Constructions were done as previously described [28] by using long PCR. Oligonucleotide primers used in this work were purchased from Integrated DNA Technologies, Skokie, Illinois. All constructs were confirmed by PCR amplification with appropriate primers (Table 1.) and DNA sequence analysis of amplicons.

**Table 1.**
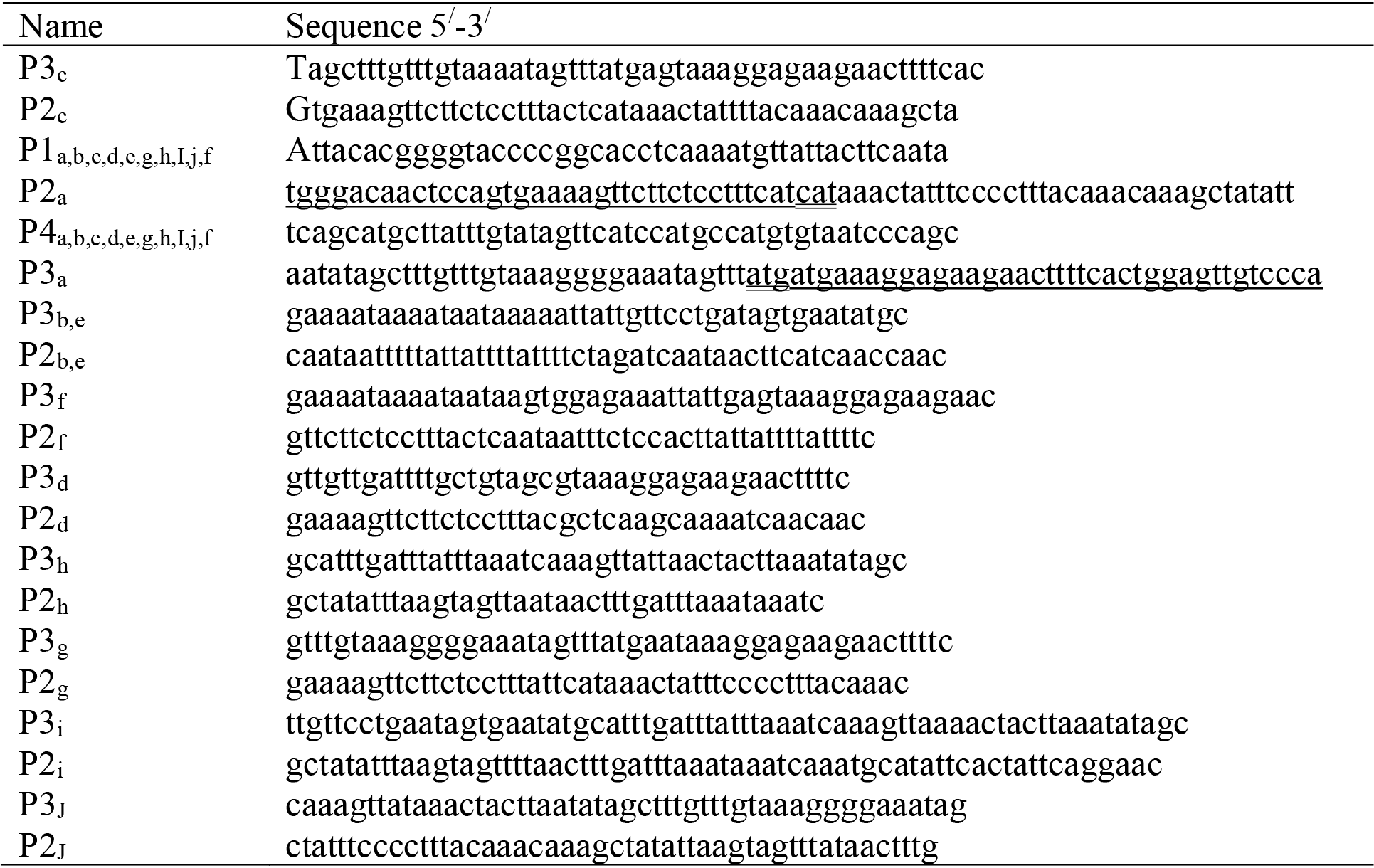
Primers used in this work.

### BmpA_L_-Gfp fusions and BmpL mutations construction

The strategy for constructing the Gfp fusions is shown in Fig.1. Different lengths of *bmpA* mRNA sequence was amplified from *B. burgdorferi* 297 total DNA with a gene-specific forward primer, P1, that annealed at least 190 bp upstream from the translational start codon in order to incorporate the native promoter and included a linker containing a specific restriction enzyme (RE) site to facilitate cloning. Primer P1 was paired with the reverse primer, P2, which included a linker that contained 25 to 30 bp *gfp*. The Gfp amplified from pCE320 [29] with primer P3, which included 25 to 30 bp of the specific BmpA sequence and primer P4, which included an in-frame stop codon and another RE site.

Deletions of SD_1_ or SD_2_, stop codons and leader sequence mutations were introduced in the primers and incorporated in the constructs by PCR. Constructs that contain both SDs and has no mutations were created first and then were used as a template to generate constructs menschen above.

Primers used to amplify the GFP and the individual BmpA sequences are listed in Table 1. To produce the fusion constructs, each BmpA fragment or mutant and GFP amplicons were mixed and amplified using P1 and P4 primers.

The PCR amplification parameters for all constructs in this work were as follows: denaturation for 2 min at 94^0^C for one cycle, followed by 38 cycles of 94°C for 10 s, 53°C for 10 s, 72°C for 2 min, and a final extension at 68°C for 5 min. The resulting PCR product was purified and cloned into pCR2.1-TOPO and subsequently electroporated into *E. coli* TOP10. Plasmid DNA from electroporants selected on Luria-Bertani agar plates with kanamycin or ampicillin (according to manufacture instruction) was purified. Then each construct was excised and subcloned into pKFSS1 [30]. DNA fragments containing cloned constructs in all structures were confirmed by DNA sequencing.

**Fig.1.**
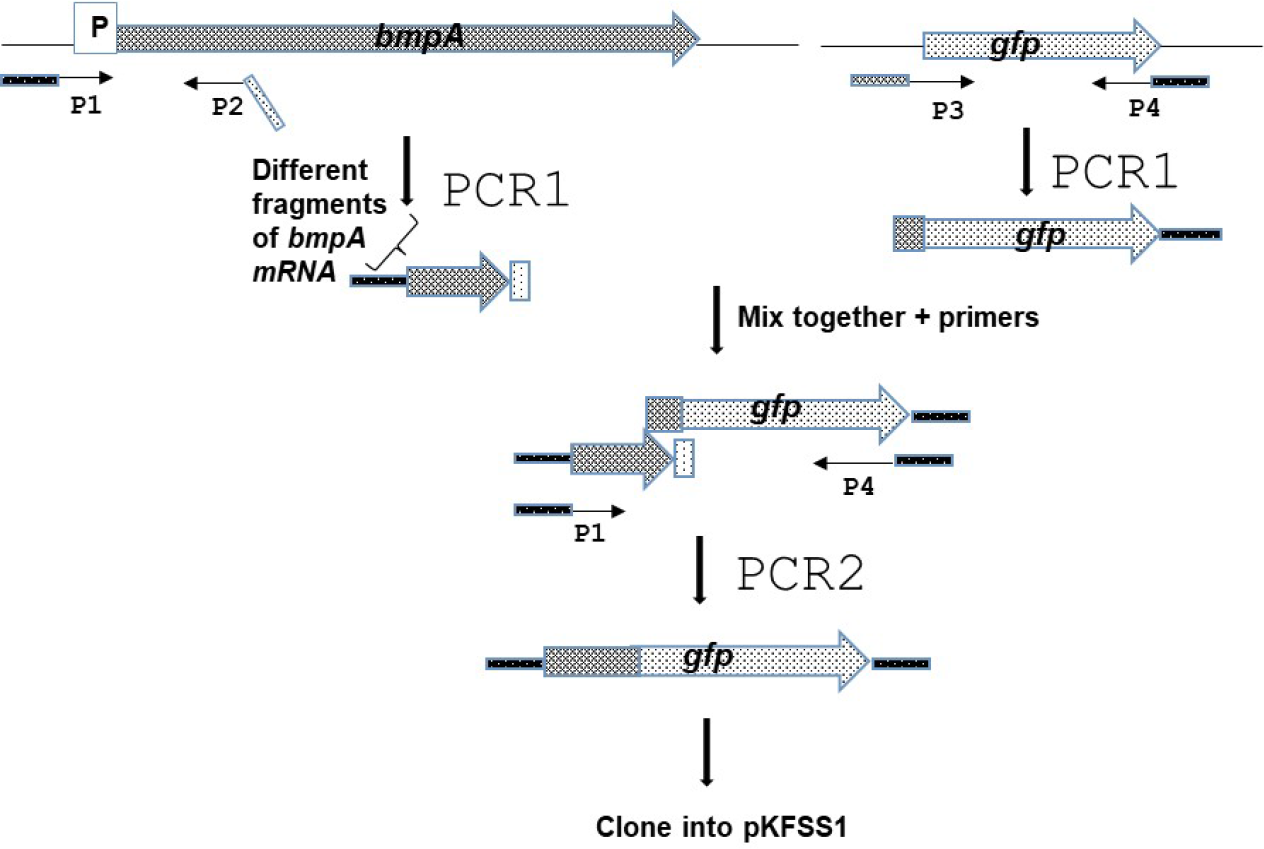
Schematic description of the construction process of the *B. burgdorferi* 297 BmpA_L_-FGP constructs. The different BmpA_L_-GFP constructs were made by truncating *bmpA*_*L*_ and *bmpA*, as well as introducing deletions of SD1, SD2 and leader mutations in the primers.

All constructions are located under native *B. burgdorferi* 297 BmpA promoter and contain different length of BmpA mRNAsequnse. Construct ***bmpA*_*L*_*::gfp*** contains mRNA *bmpA* sequence from −190 base pair (bp) to *bmpA* starting codon (-<u>AUG</u>-) and *gfp* under this start codon. Constructs ***bmpA*_*L*_(ΔSD_1_)::*gfp*** and ***bmpA*_*L*_(ΔSD_1_)::*gfp*** differ from first one by deletion of SD_1_ (-<u>GTGGAG</u>-) and SD_2_ (-<u>AGGGGA</u>-), respectively. In constructs ***bmpA*_*L*_(33bp*bmpA*)::*gfp*,** *gfp* starts after 33 bp of *bmpA* gene respectively, and in ***bmpA*_*L*_ SD_1_::*gfp*** contains *gfp* starts after *bmpA*_*L*_ start codon -<u>UUG</u>-. In the ***bmpA*_*L* stop_::*gfp*,** *gfp* is fused after leader peptide stop codon.

### The *B. burgdorferi* electroporation

*B. burgdorferi* 297 at mid-log phase (1-2 × 10^7^ cells/ml) was electroporated with 10 to 30 μg of recombinant plasmid DNA. After overnight recovery, cells were diluted to 10^7^ cells/ml and distributed into 96 micro-well plates (Corning Incorporated, Corning, N.Y.) containing BSK-H media with 70-100 μg/ml of streptomycin for selection of clones containing recombinant plasmid. After 10-15 days DNA of *B. burgdorferi* cells growing in these microwells was checked for the presence of the plasmid by fluorescence and by PCR. The DNA of streptomycin resistant colonies was extracted using High Pure PCR Template Preparation Kit (Roche Diagnostics Corporation, Indianapolis, IN) and analyzed by PCR for the presence of the appropriate construct with specific primers (Table 1.).

### Detection of GFP and BmpA by immunoblotting

*E. coli* DH5α and *B. burgdorferi* 297 total proteins were extracted from 1-2×10^7^ cells/ml by lysing them in Laemmle buffer. Protein lysates were analyzed by SDS-PAGE followed by silver stain or immunoblotting using rabbit anti- GFP (Invitrogen, Eugene, Oregon, USA) or anti-BmpA polyclonal antibody. Immunoblots were developed using ECF Western Blotting Kit according to the manufacturer’s instructions (Amersham Biosciences, Piscataway, N.J.), and detected using a Storm 860 PhosphorImager and ImageQuaNT software (Molecular Dynamics, Sunnyvale, CA).

### Flow cytometry analysis

Aliquots from three independent experiments, containing *E. coli* at 0D_06_=08 and 1×10^8^ *B. burgdorferi* B31 and its derivatives containing GFP in pKFSS1 or TOPO were washed with PBS and analyzed on a FACS scan flow cytometer (Becton Dickinson, Mountain Lake, Calif.) using CELLQUEST 3.2 (Becton Dickinson).

### Microscopic analysis

Cultures of E. *coli* and *B. burgdorferi* 279 that contain different constructs (10^6^ cells/ml) were examined by fluorescence microscopy to detect the fluorescence.

## Results

### Analyze the *bmpA* gene

The transcription start site of the *bmpA* gene is set at −105bp position relative to its translational initiation codon and is located within the coding sequence of the *bmpC* gene (60 bases upstream from the *bmpC* stop codon).

There are several palindromic sequences in the *bmpA*_*L*_ present, suggesting that *bmpA*_*L*_ can also form complicated secondary structures and two purine reach regions that can serve as a ribosome binding site (RBS) [31–33]. Moreover, the ORFs of a leader peptide and BmpA can form two different frames and contain a stop codon for a BmpA_L_ that can overlap with the start codon of BmpA (Fig. 2A) suggesting that these two proteins can be co-expressed and coregulated. Introducing mutations to the palindromes, in the way that they change the mRNA secondary structure but not affect the amino acid sequence, does not affect BmpA_L_ and BmpA expression (data are not shown).

**Fig.2.**
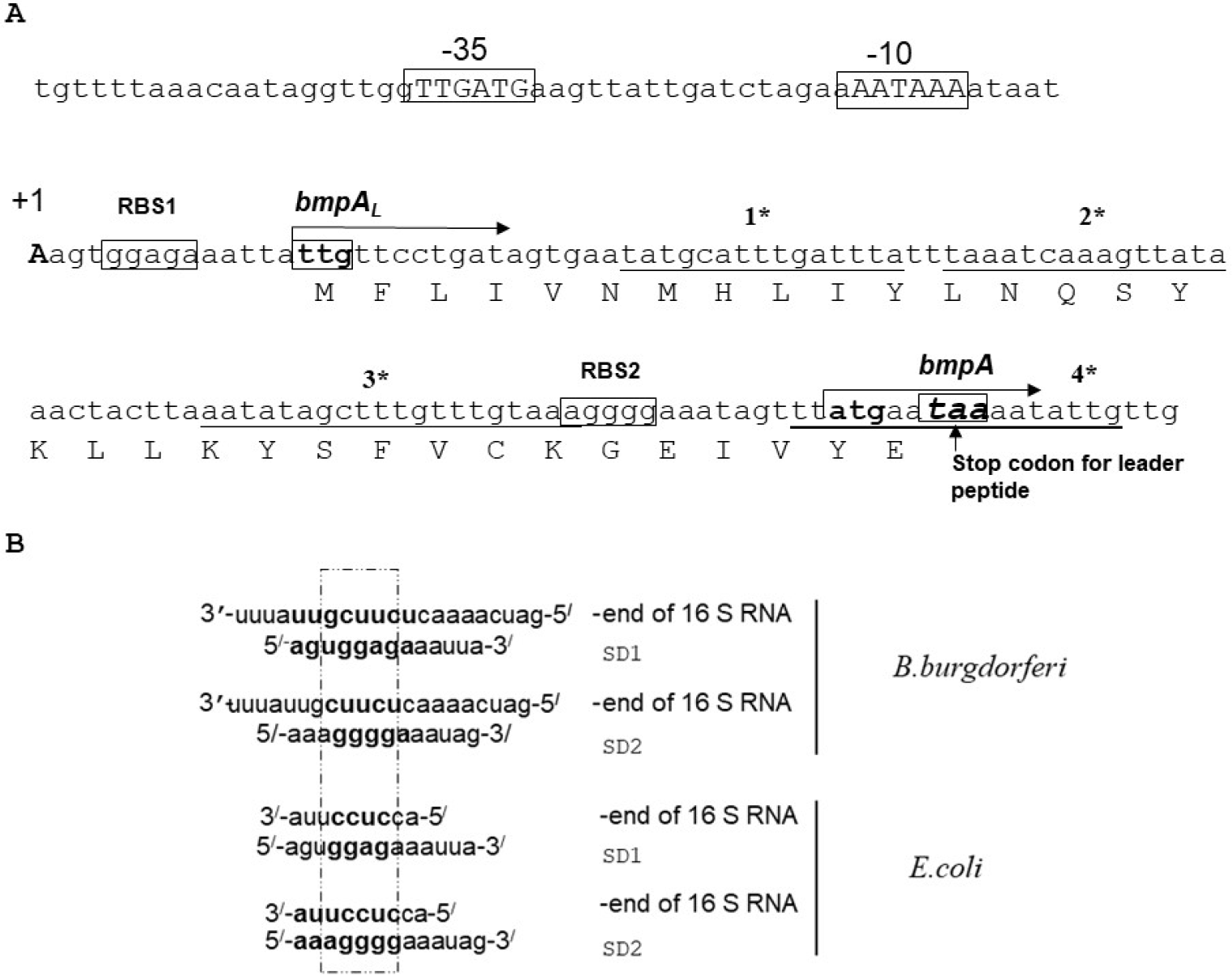
The nucleotide sequence of the 5’-flanking regions of the *bmpA* gene. **A**. Transcription initiation site is indicated as +1. Putative *bmpA* leader peptide (*bmpA*_*L*_) is also shown. Translational initiation codons for *bmpA*_*L*_ and *bmpA* are boxed; stop codon for leader peptide is indicated by the arrow. The invert rapids are indicated by the star. **B**. Alignment of the *B.burgdorferi* SD sequences with 3^/^ end of 16S rRNA. The two Shine – Dalgarno (SD) sequences found in the *bmpA* mRNA can base pair with the 3^/^ end of the 16S rRNA.

The SD_1_ sequence is -<u>GGAG</u>- with a spacing between this SD_1_ and the initiation codon <u>-UUG-</u> of 10 bp as counted from the first G in SD_1_ (Fig. 2A). The spacing between SD_2_ (-<u>AGGGGA</u>-) and the initiation codon -<u>AUG</u>- is 12 bp counted from the second G at position 3 in SD_2_ (Fig. 2A). Both SDs in *bmpA* mRNA can pair with the 3’ end of the 16S rRNA (Fig. 2B) [31] suggesting that both SDs can be functional in *B. burgdorferi*.

### Determination of RBS for *bmpA*

Careful sequence analysis of the *bmpA* gene shows the presence of two potential SDs. The first SD_1_ (-<u>GGAG</u>-) starts at nucleotide position +4 counting from transcription start codon and is close to an alternative start codon -<u>UUG</u>-. The sequence (-<u>GGAG</u>-) is classical SD sequence for many species of bacteria and was found approximately in 43% genes of *B. burgdorferi* when searched in PubMed database.

The distance between (-<u>GGAG</u>-) and the translation start codon for *bmpA* is 100 bp. The effects of SD spacing, distance between SD and the initiation codon, variation in SD sequences and the effects of other alternative translational start sites are well studied. The excessively long, or short spacing between the SD and the initiation codon may abolish or limit efficient translation initiation [34, 35].

The second SD (-<u>AGGGGA</u>-) is located at nucleotide position 90 counting from +1 and 14 nucleotides from the described translation start codon <u>-AUG-</u> for *bmpA*. Sequence ‒<u>AGGGGA</u>- is less common as an SD and does not appear as an SD sequence in the database for *B. burgdorferi*. Moreover, two of the predicted SDs can pare with 16S rRNA (Fig. 2B).

To verify -<u>GGAG</u>- or -<u>AGGGGA</u>- is an SD for BmpA we made several constructs that differ only in *bmpA*_*L*_. One, of these, contain the *bmpA* promotor *bmpA*_*L*_ and the translation start codon -<u>AUG</u>- of *bmpA* fused to *gfp* (*bmpA*_*L*_*::gfp*). A second differs from the first one only by a deletion in the SD_2_ (-<u>AGGGGA</u>-) sequence ((*bmpA*_*L*_(ΔSD_2_)::*gfp*)), and the third construct contains deletion of the SD_1_ (-<u>GGAG</u>-) sequence (*bmpA*_*L*_(ΔSD_1_)::*gfp*). Expression of *gfp* was studied in both *E. coli* which served as a model microorganism as well as in *B.burgdorferi* strain 297.

The results of flow cytometric analysis are presented in Fig.3. In the plasmid that harbored *bmpA*_*L*_,(ΔSD_2_)::*gfp* (Fig.3. line 2) fluorescence in *E. coli* and *B. burgdorferi* strains were not detected. In opposite, deletion of SD_1_ did not abolish the GFP expression (Fig. 3. line 3), and in *E. coli* GFP expression was at the same level as in construct *bmpA*_*L*_*::gfp* that contains both SD sites. At the same time, in *B.burgdirferi* 297 GFP expression was approximately twice higher comper to GFP expression from construct *bmpA*_*L*_*::gfp*. This data suggests that -<u>AGGGGA-</u> is indeed an SD site of BmpA. Moreover, the facts that both SDs (-<u>GGAG</u>- and -<u>AGGGGA-</u>) can pair with 3^/^ end of 16S rRNA of *B. burgdorferi* and may form the translation initiation region (SD, initiator codon, and a spacer region) suggest that both SDs can be active (Fig.2B, 2 B).

### Detection of the leader peptide

To verify that SD_1_ (-<u>GGAG</u>-) is active and can form translation initiation region (TIR) together with -<u>UUG</u>-, we constructed plasmid in which *gfp* was fused with the first start codon -<u>UUG</u>- after predicted SD_1_- (-<u>GGAG</u>-) (Fig.3. line 4). This plasmid allowed GFP production from the start codon for leader peptide under the control of native *bmpA* promoter (P_*bmpA*_) only if -<u>GGAG</u>- plays the role as an SD sate and -<u>UUG</u>- as a start codon. Expression of GFP from this construct was studied in *E. coli* and *B. burgdorferi* and compared with expression of GFP from the plasmid that contains both SDs (Fig.3. line 1).

Flow cytometry analysis showed expression of GFP in *E. coli* and *B. burgdorferi* from the plasmid that harbored *gfp* fused in frame to a start codon of the leader peptide. Expression of GFP from this construct was also detected by western blotting (data not shown). This result indicates that the BmpA_L_ is translated from -<u>UUG</u>- start codon using -<u>GGAG</u>- as SD_1_. The low-level expression may be explained by rearly used start codon -<u>UUG</u>-.

**Fig. 3.**
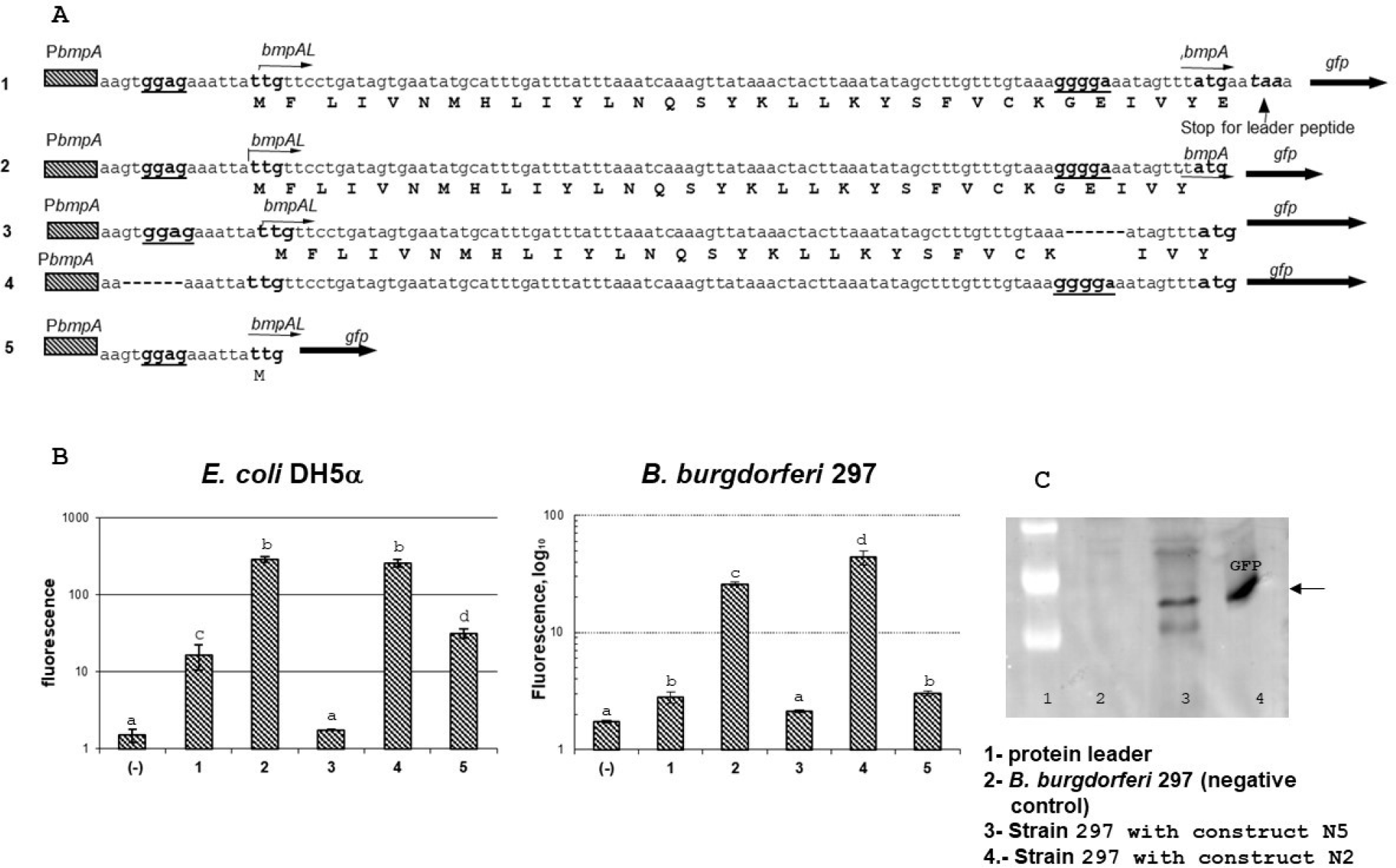
Expression of GFP in recombinant strains. **A**. Sequences of different constructs fused to *gfp*. **1**. The *bmpA*_*Lstop::*_*gfp* construct, that contains *bmpA* promoter *bmpA* leader from +1 to the BmpA stop codon and *gfp* fused in frame with BmpA start codon. **2**. The *bmpA*_*L*_::*gfp* construct, differ from construct one by fusion *gfp* directly to the bmpA start codon -ATG-. **3.** The *bmpA*_*L*_(ΔSD_2_)::*gfp* construct, differ from the second construct by deletion of SD_2_ (-<u>AGGGGA</u>-). **4.** The *bmpA*_*L*_(ΔSD_1_)::*gfp* construct, differ from the second construct by deletion of SD_1_ (-<u>TGGAGA</u>-). **5**. The *bmpA*_*L*_ SD_1_::*gfp*, contains P_*bmpA*_ and *bmpA*_*L*_ from +1 to -<u>UUG</u>- (start codon for leader peptide) fused in frame to *gfp.* **B**. Expression of GFP detected by flow cytometry in *E. coli* and *B.burgdorferi* recombinant strains. Level of GFP expression from the constructs: **1**) *bmpA*_*Lstop::*_*gfp;* **2)** *bmpA*_*L*_::*gfp*; **3)** *bmpA*_*L*_(ΔSD_2_)::*gfp*; **4)** *bmpA*_*L*_(ΔSD_1_)::*gfp*; **5)** *bmpA*_*L*_ SD_1_::*gfp*. Statistical analysis was conducted using 1-way ANOVA followed by Tukey’s post hoc test for pairwise comparisons. Data are mean ± SD of 3 replicates; columns with the same letters are not significantly different (p < 0.05). **C**. Representative Immunoblot for detection of GFP expression in recombinant strains *E. coli* and *B. burgdorferi*.

### Expression of leader peptide inhibits *bmpA* gene expression

To detect that leader peptide expression has any effect on BmpA expression we created a construct that contains *PbmpA, bmpA*_*L*_, and 33 bp of *bmpA* fused in frame with *gfp* protein. The second construct was created from a first one by deletion of sequence -<u>GGAG</u>- that corresponds to SD_1_ (*bmpA*_*L*_(ΔSD_1_)33bp*bmpA::gfp*). Expression of GFP was significantly higher in the *E. coli* and *B. burgdorferi* strains that contain SD_1_ deletion (*bmpA*_*L*_(ΔSD_1_)33bp*bmpAgfp*) compared to strains that contain both SDs *bmpA*_*L*_ 33bp*bmpAgfp* construct (Fig. 4. A, B).

*B. burgdorferi bmpA* monocistronic message contains two SDs. The fact that SD_2_ is active even when SD_1_ is deleted suggests that SD_2_ is not translationally coupled to SD_1_ by secondary structure, moreover elevated level of expression in the case where SD_1_ was removed compare to the construct that contains both SDs suggest that translation of BmpA_L_ inhibits BmpA translation (Fig. 3. and Fig. 4). This effect was not detected in E. *coli* strains and can be explained by stronger pairing of 16sRNA with SD_2_ compare to SD_1_ (Fig. 3B).

To verify that stop codon for leader peptide plays any role in regulation of the upstream located gene, we created a construct that contains entire *bmpA*_*L*_ including stop codon and GFP fused in frame with -<u>AUG</u>- of the *bmpA* gene (*bmpA*_*L stop*_::*gfp*). The expression of GFP was detected by flow cytometry (Fig. 3). Presence of stop codon significantly inhibited *gfp* translation, compare to construct were gfp was fused directly to a start codon of bmpA. Moreover, as we expected, according to ribosome pairing with SD in *E. coli* and *B.burgdorferi*, the effect was more noticeable in B. *burgdorferi* compare to *E.coli*, suggesting that stop codon of BmpA_L_ plays significant role in the expression of *bmpA* gene.

We also introduced stop codon inside of the leader peptide (Fig. 4. construct 3). Western blot analysis shows expression of GFP in this construct only in *E. coli*, and not in *B.burgdorferi* (Fig. 4B).

Thus, our results demonstrate that SD_2_ is not translationally coupled to SD_1_ by secondary structure, translation from SD_2_ does not require SD_1_, and translation from SD_1_ inhibits translation from SD_2_.

**Fig.4.**
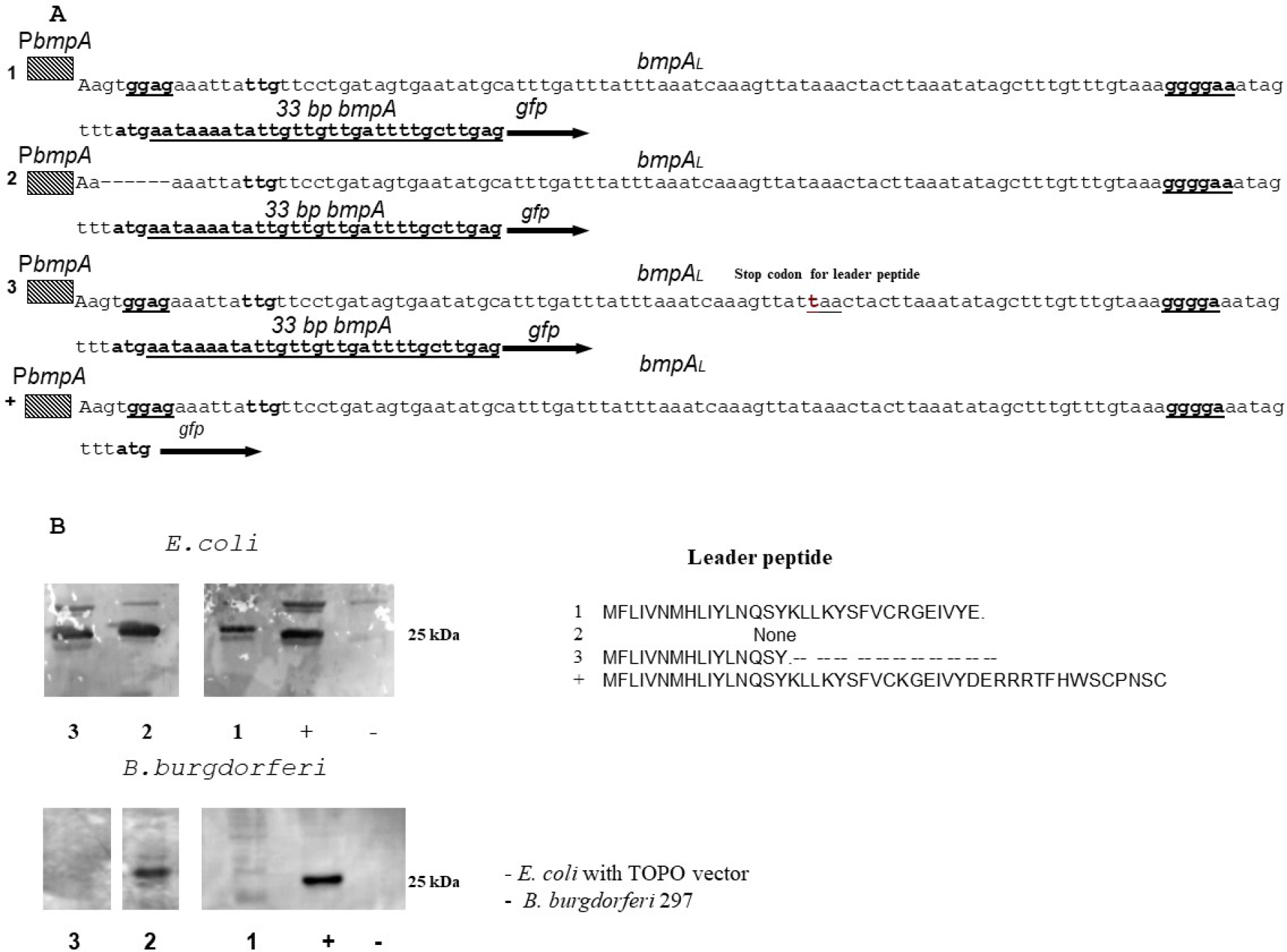
Leader peptide expression inhibits expression of BmpA. **A.** Sequence of different constructs fused in frame with *gfp*. **1.** *bmpA*_*L*_33bp*bmpA::gfp* construct, that contain *bmpA* promoter, *bmpA*_*L*_, 33bp of *bmpA* fused in frame with *gfp*. **2**. The *bmpA*_*L*_(ΔSD_1_)33bp*bmpA::gfp* construct, differs from construct one by deletion of SD_1_ (-<u>TGGAGA</u>-). **3.** The *bmpA*_*L***ochre17**_n33bp*bmpA::gfp* construct, that contains *bmpA* promoter *bmpA* leader with the stop codon in position 17, 33bp of *bmpA* fused in frame with *gfp*.**4**. The *bmpA*_L::*gfp*_ construct, contains *bmpA* promoter *bmpA*_*L*_ leader from +1 to the BmpA start codon that fused in frame with *gfp*. The *bmpA*_*L::*_*gfp* was used as positive control **B.** Western blot analyses expression GFP from different construct is described above.

### Comparison of *bmpA*_*L*_

*B.burgdorferi* ORF for *bmpA*_*L*_ encodes 32 amino acids leader peptide with molecular weight 3882.67 Daltons. It contains three strongly basic (+) amino acids (K, R), two strongly acidic (−) amino acids (D, E), fourteen hydrophobic amino acids (A, I, L, F, W, V), and ten polar amino acids (N,C,Q,S,T,Y). The Isoelectric Point of this peptide is 8.178, 1.044 Charge at Ph 7.0. Nucleotide sequence for *bmpA*_*L*_ contains % A+T = 76.77% C+G = 23.23% where % A = 36.36; % G = 16.16; % T = 40.40; % C = 7.07.

The BmpA_L_ amino acid sequence shows strong similarity to other species of *Borrelia* leader peptide but we do not find homology to another bacterial leader peptides. First 19 amino acids are strong conservative (Fig. 5). Inside of this conservative region located 5 leu codons and 3 of them rarely used in *Borrelia*, suggesting that they can play a regulatory role.

**Fig.5.**
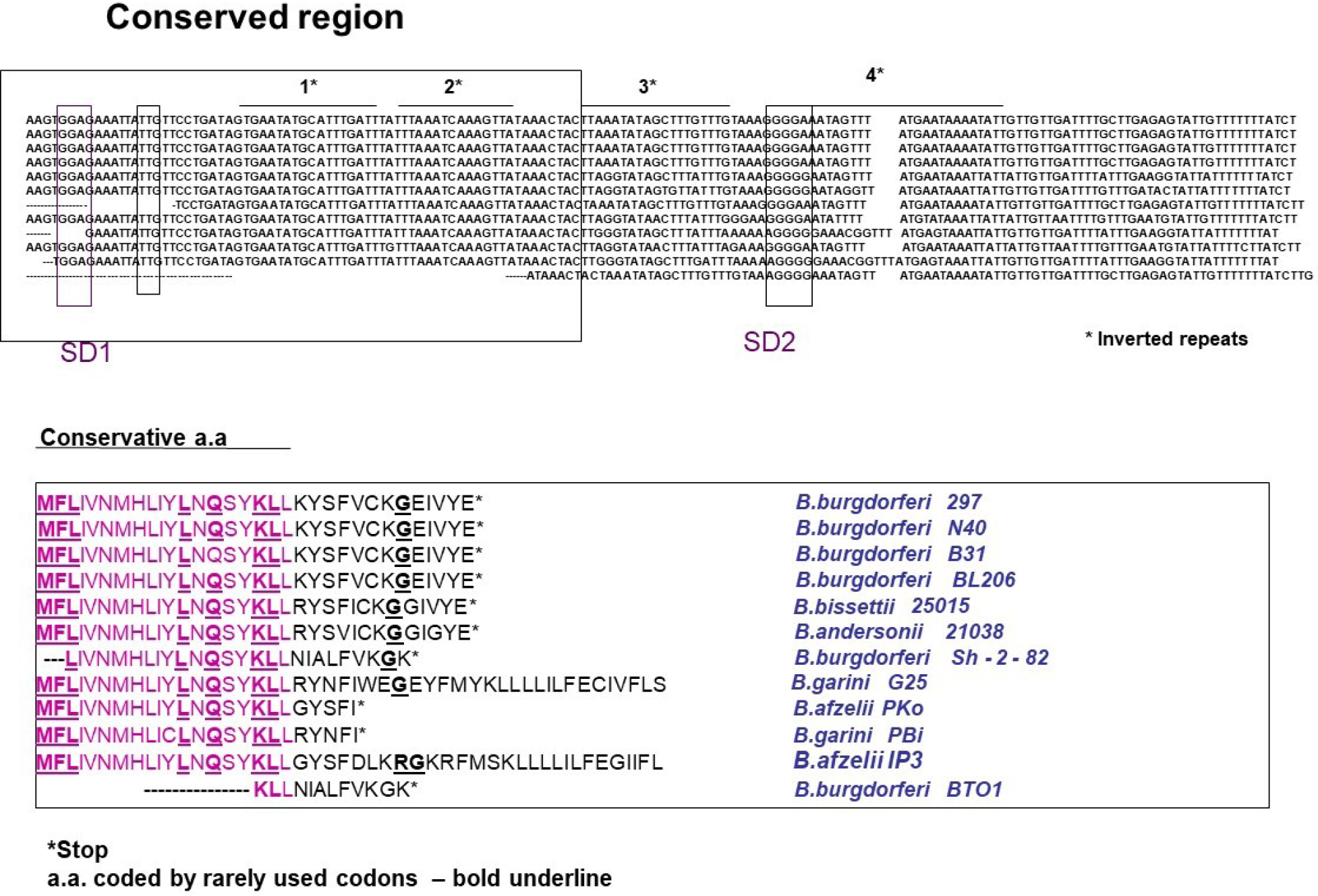
Sequence alignment of the leader peptides from different *Borrelia* species. Sequences alignments were done using DNA star. Conservative amino acids are boxed in the DNA sequence and red in amino acids sequence. Stop codon is indicated by a star and rarely used codons-bold underline.

Moreover, BmpA_L_ amino acid sequence also has significant differences between *Borrelia* species. *B. burgdorferi* strains 297, N40, B31, BL206 have conservative 32 amino acids leader peptide with stop codon located two nucleotides after start codon for *bmpA*. The *B. bissettii* 25015 and *B. andersonii* 21038 also have 32 bp leader peptide but different in amino acids in the variable region.

*B. burgdorferi* SH-2-82 and *B. burgdorferi* BTO1 have shorter-28-amino acids peptide. *B. afzelii* PKO and *B. Garini* Pbi have 24 leader peptide, and *B. afzelii* IP3 or *B. garini* G25 has leader peptide in frame with *bmpA*. This data may suggest strain-depenmdent differences in *bmpA* regulation and expression. For example, *B. afzelii* IP3 or *B. garini* G25 can use the SD_1_ or SD_2_ for expression of BmpA. It can contain two BmpA products with and without the leader sequence. At the same tame expression *bmpA* in strains *B. burgdorferi* SH-2-82, *B. burgdorferi* BTO1, *B. afzelii* PKO and *B. Garini* Pbi can be reinitiated from ORF started from SD_1_. It is not clear if these phenomena have a biological importance.

### The variable sequence of the leader peptide can be important for the inhibition of *bmpA* translation

Two frameshifting mutations were introduced to the *bmpA*_*L*_ to examine role of conservative and variable parts of the *bmpA*_*L*_ on *bmpA* translation. The first two-point mutations alter amino acid sequence between *bmpA*_*L*_ 4aa and 16 aa. Expression of GFP in resulting mutant does not differ from wild-type *bmpA*_*L*_. At the same tame a change in amino acids sequence of variable part (deletion A in codon 20) significantly increases GFP expression (Fig. 6). Moreover, in this case, BmpA_L_ is shorter similarly to *B. burgdorferi* Sh-2-82 where stop codon located immediately before SD_2_, indicating that translation at SD_2_ can reinitiate from ORF started at SD_1_.

**Fig. 6.**
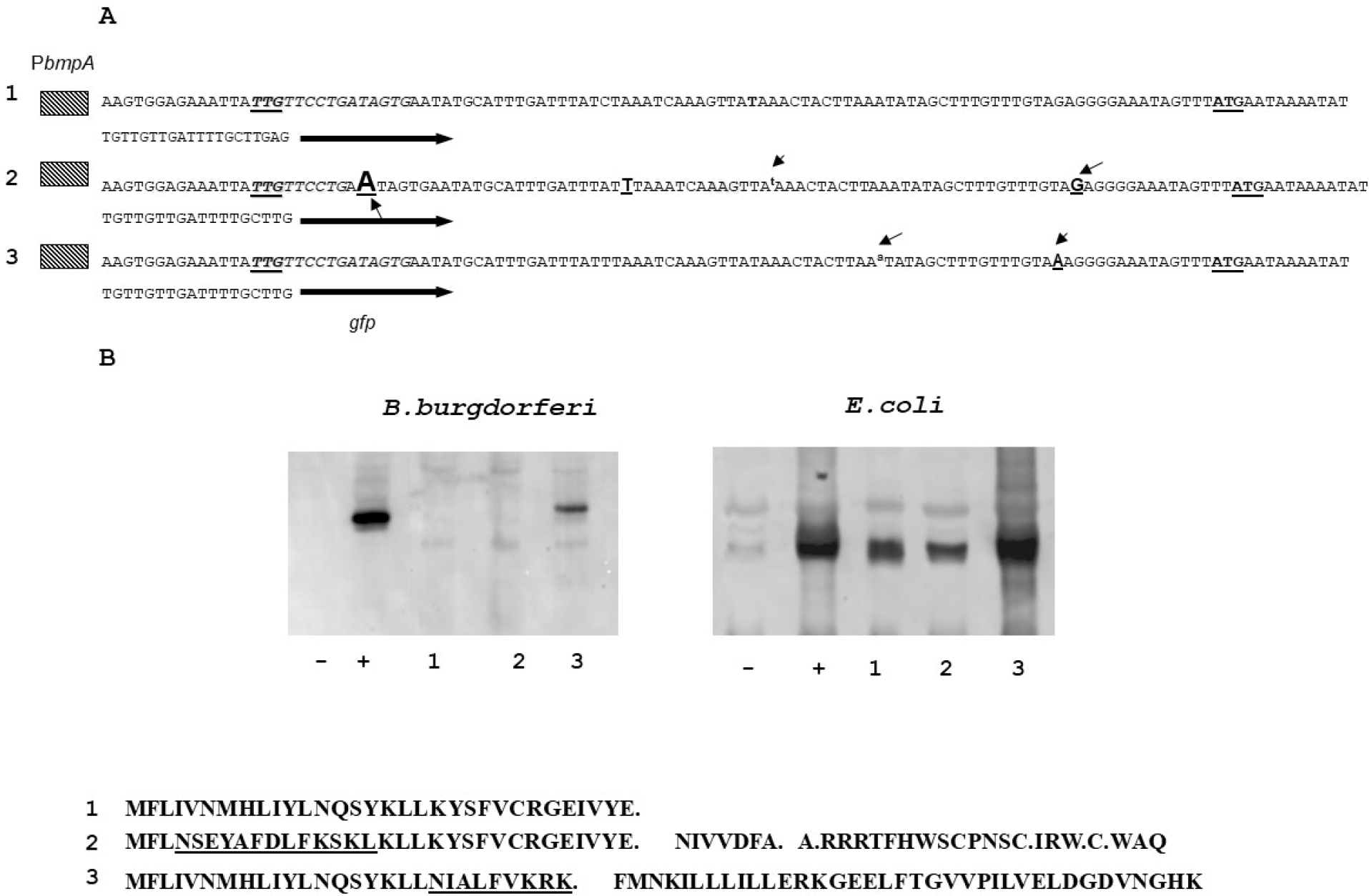
The GFP expression from recombinant strains containing constructs with mutations in conservative and variable part of *bmpA*_*L*_. **A.** Sequence of different constructs fused to *gfp* that was used for this study: **1.** *bmpA*_*L*_33bp*bmpAgfp* construct that contains *bmpA* promoter, *bmpA*_*L*_, 33bp of *bmpA* and *gfp* fused in frame to *bmpA*. **2**. The *bmpA*_*L*mutatedconserve_33bp*bmpA*::*gfp* construct difer from the first one by insertion of <u>-A-</u> in position 25 and deletion of <u>-T-</u> in position 62. **3**. The *bmpA*_*L*mutatedvariable_33bp*bmpA*::*gfp* construct, contains deletion of <u>-A-</u> in position 74. **B.** Expression of GFP detected by western blotting in *E.coli* and *B. burgdorferi*. Recombinant strains that contain: **1.** *bmpAl*33bp*bmpAgfp;* **2**. *bmpA*_*L* mutatedconserve_::*gfp;* **3**. *bmpA*_*L*mutated variable_::*gfp*.

### Working model

Based on results, described above we proposed a model similar to the models described for *E. coli* [36, 37] and eukaryotic protein-encoding genes that contain upstream ORFs [38] (Fig. 7). In strain 297, ribosome proceeds starting from an SD_1_, it then overrides the SD_2_, so SD_2_ site becomes silent (A). In *B. afzeliiPKo, B. garini PBi, B. burgdorferi BTO1* the ribosome can reinitiate translation from ORF starting from SD_1_ (B). In *B. garini G25, B. afzelii IP3* SD_2_ leader peptide fused in frame with the *bmpA* gene. In this case, two products are possible. First one contains BmpA protein together with leader peptide, another one only BmpA. To test this hypothesis, we performd the western blotting on *B. burgdorferi* 297, N40 and *B. afzelii* IP3 (Fig, 8). Data indicates that BmpA of *B. afzelii* is slightly larger compared with BmpA of *B. burgdorferi* 297 and N40, confirming our hypotheses.

**Fig. 7.**
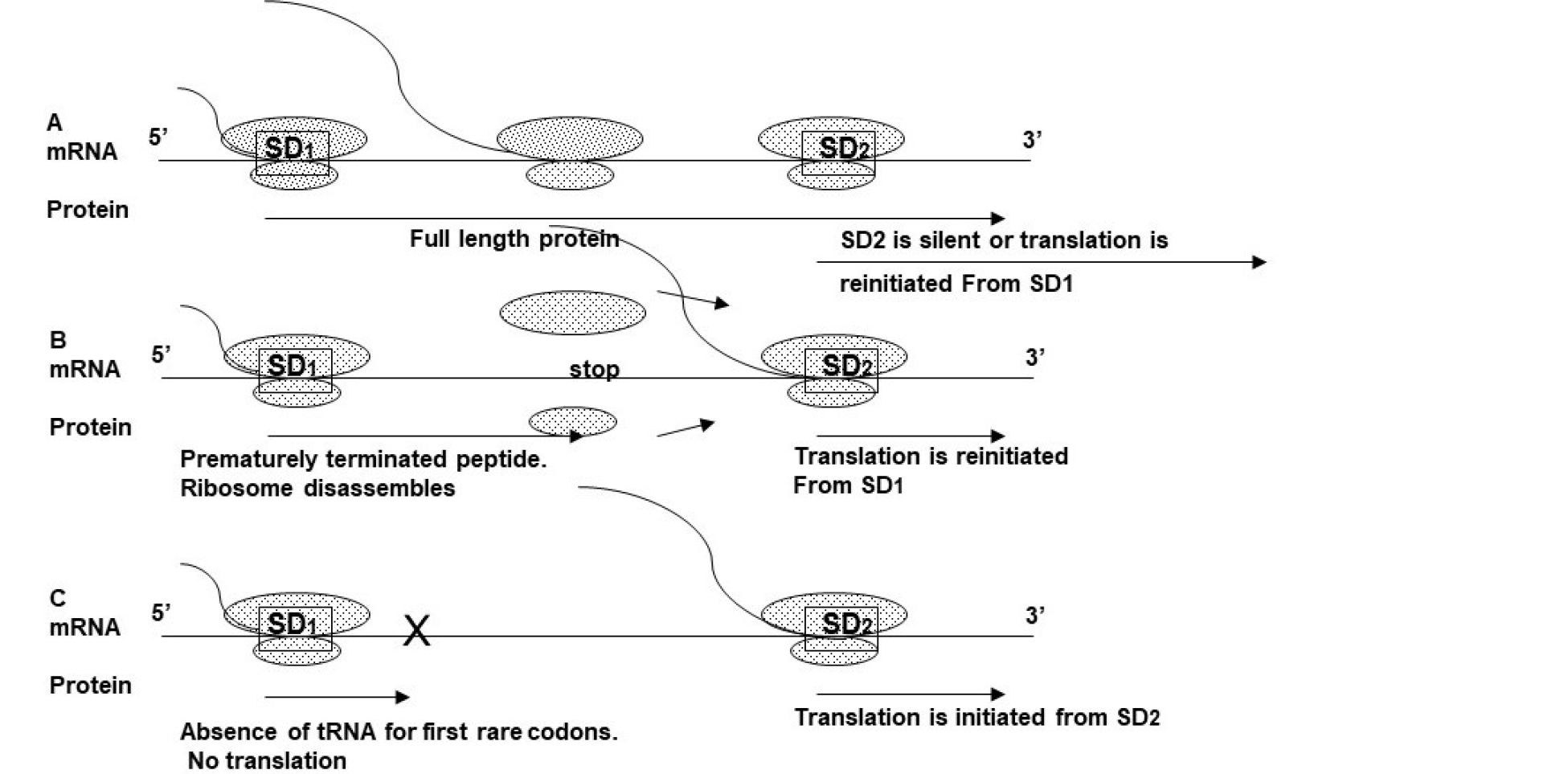
Working model. A. In 297 wild-type ribosomes that began translation from SD_1_ pausing at -<u>GGG</u>- and at stop codon, preventing second ribosome polymerization at SD_2_. Ribosomes also can re-initiate translation from SD_1_. B. In strains with a stop codon before SD_2_ the translation from SD1 is terminated, and ribosome can reinitiate translation at SD_2_. C. Translation initiation from SD_2_ when SD_1_ is abolished.

**Fig.8.**
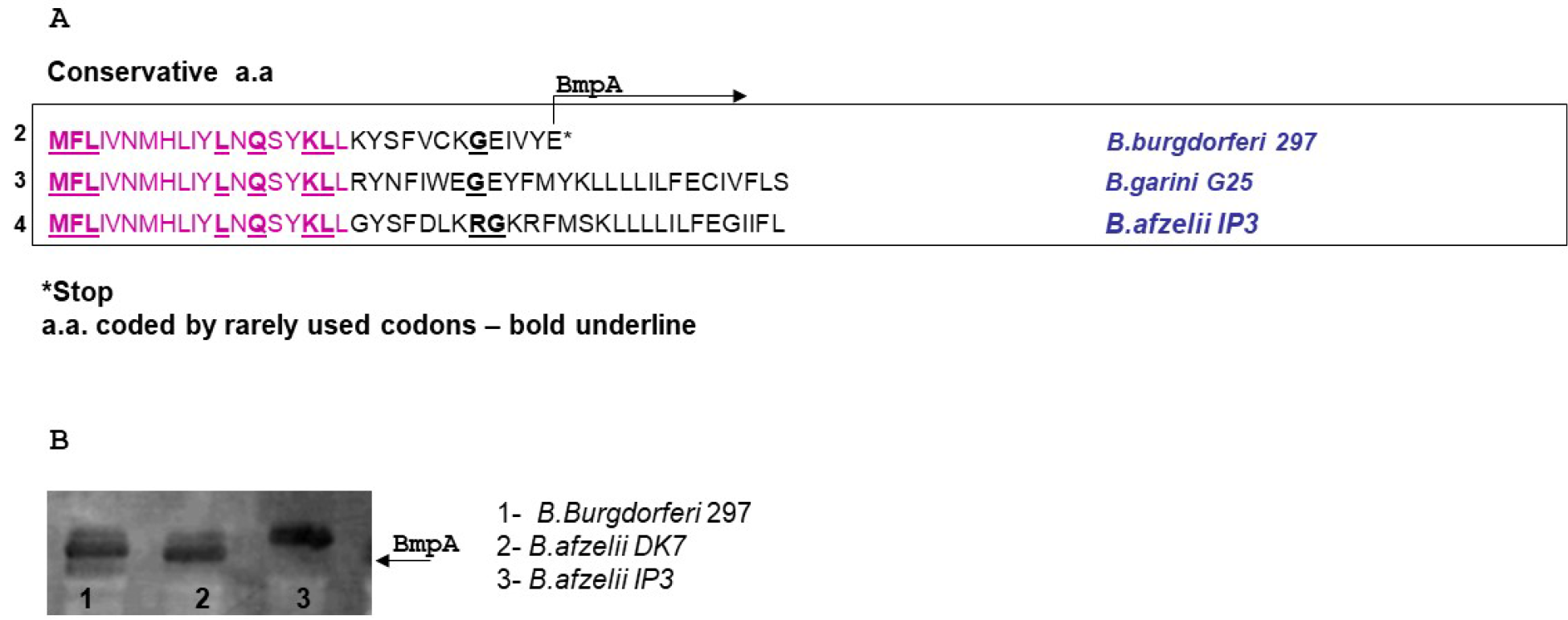
Expression of BmpA from different *B. burgdorferi* strains. A. The BmpA_L_ amino acid sequences of *B. burgdorferi* 297 (2), *B. garini* G25 (3), *B. afzelii* IP3 (4). Amino acids coded by rarely used codons, bold underline. B. Immunoblotting of BmpA expression from different B. *burgdorferei* strains.

The lower level of translation from constructs that contain both SD_s_ compare wit the same constructs that contain only SD_2_ can be explained by *bmpA*_*L*_ sequence. The first four codons of leader peptide are rare for *B. burgdorferi*, suggesting that ribosome may translate this region slower. Another fact, that leader peptide inside of conservative region contains five leucine (Leu), codons and three of them are rarely used for *B. burgdorferi*, suggest that they can be involved in regulation of leader peptide expression. Moreover, another rare codon <u>-GGG-</u> in *B. burgdorferi* bmpA_L_ is located in SD_2_ region and can slow down ribosome movement, covering SD_2_ and inhibiting polymerization second ribosome and translation from SD_2_. We also do not exclude that additional RNA binding factors or secondary structure of 5’RNA may also play a role in the regulation of leader peptide expression.

## Discussion

Post-transcription regulation of gene expression is a key mechanism by which cells and organisms can rapidly change their gene expression in response to internal or external stimuli. Expression of all genes is regulated at multiple post-transcriptional steps including mRNA stability, and translation of mRNA. Translational regulation at the initiation step can be mediated via different *cis*-acting elements present in the 5′RNA leader sequence, such as the secondary structure of the 5’ RNA and upstream open reading frames (uORFs). The uORFs can significantly change protein expression levels by interfering with the efficiency of translation initiation of the downstream ORF [38, 39], indicating that they can control protein synthesis.

Taken together, this data indicates presence of two translation initiation regions in *bmpA* mRNA. The SD_2_ is active only when SD_1_ is silent. 16S rRNA in the 30S ribosomal subunits plays a significant role in selecting the translational start site [32, 33]. In most mRNA 4 or 5 bp SD interaction is strong enough to mediate efficient translation [40]. A stronger than regular SD interaction does help, however, when the start codon is not -<u>AUG</u>-, or when the initiation site is masked by secondary structure [41]. A/U ‒rich initiation site that forms unstable secondary structure might require no SD interaction at all [42].

Stronger base pairing of SD_1_ sequence with 16S rRNA (8 bp) compare with SD_2_ (5 bp) in *B. burgdorferi* (Fig. 2B), suggests that ribosome should polymerize more efficiently in SD_1_. In *E. coli* the pairing 16S rRNA to SD_1_ and SD_2_ is opposite 4 and 6 bp, correspondingly. This fact indicates that in *E. coli* the ribosome is polymerized more efficiently in SD_2_. The -<u>UUG</u>- uses about 3% of the start codons in *E. coli* and it is also rare start codon for a *B. burgdorferi*. The - <u>AUG</u>- start codon is preferred via pairing with the anticodon (-<u>UAC</u>-) in fMet-tRNA. Weaker pairing is part of the reason for less efficient translation when -<u>GUG</u>- or -<u>UUG</u>- is used as a start codon. In *E. coli* translation from these codons are 8 times less efficient than from -<u>AUG</u>- [43]. Therefore, even when SD_1_ have more bases pairing with 16S rRNA in *B. burgdorferi* the translation of the BmpA_L_ is reduced by using -<u>UUG</u>- as the start codon.

Many bacterial genes are parts of polycistronic operons [44–47]. Translation coupling and re-initiation are important for the expression of functionally related proteins from polycistronic operons [48, 49]. The cistrons of some translationally coupled messages do not have an independent SD, and in this case, the stop codon of upstream cistron and the initiator codon of the downstream cistron overlap. A functional ribosome in stop codon for the first peptide reinitiates translation of downstream cistron instead of getting disassembled. A defect in translation from the first cistron abolishes the translation from the downstream cistron [50]. *B. burgdorferi bmpA* is transcribed as a monocistronic mRNA that contains two SDs, and as polycistrons with *bmpCbmpA* and *bmpAbmpB*. Removing the SD_1_ from this mRNA increases translation of GFP or BmpA in *E. coli* and *B. burgdorferi* suggesting that SD_2_ is not translationally coupled to SD_1_ by secondary structure (Fig. 3, 4, 6).

The size of the ribosome is 25nm, indicating that one ribosome may occupy the space approximately 10-20 aa [51]. When ribosome begins translation from SD_1_, another ribosome has enough space for polymerization and translation from SD_2_. But in the absence of translation termination, the ribosomes from SD_1_ move forward and cover the area necessary for ribosome polymerization at SD_2_.

The rate at which elongating ribosome translates through ORF is codon-specific and in *E. coli* differ from 5-21 codons per second. The ribosome can stall during translation elongation in rare or termination codons, creating a blockade to addition ribosome. The ribosome stalling is also involved in positive regulation of translation [51–53].

The first four amino acids in the leader are rare codons and may regulate translation efficiently from SD_1_. When ribosome occupies two leader codons for Leu, the ribosome progressively encroaches on the space needed for a second ribosome to initiate at SD_2_. The fact, that *bmpA*_*L*_ contains codons that are used less efficiently in *Borrelia* may also be the reason for a slower translation of *bmpA*_*L*_. Finally, the rare codons for Cys (-<u>UGU</u>-) in the variable part of *bmpA*_*L*_ and Gly (-<u>GGG</u>-) inside of SD_2_ as well as the *bmpA*_*L*_ stop codon are the reasons of slower ribosome translation from SD_1_ that interfere with the second ribosome polymerization at SD_2_ [53].

When native *bmpA*_*L*_ together with a stop codon was present in the derivative plasmids, GFP expression was significantly lower in *E. coli* and not detected or detected on the low level in *B. burgdorferi* compare with similar constructs that do not have the stop codon (Fig. 3). GFP translation from the construct (Fig. 6) in which the stop codon is located before the SD_2_, was significantly higher compared with translation from construct containing native *bmpA*_*L*_ (Fig. 7). Moreover, the aa sequence of this construct reminds the sequence of a *B. burgdorferi* Sh-2-82 and *B. burgdorferi* BTO1. This data suggests that translation from SD_2_ can happen *de novo* or by re-initiation from SD_1_.

Lyme disease patients, having Borrelia burgdorferi infection, show variety of clinical evidences from asymptomatic infection to chronic arthritis. The most common clinical sign of infection is an erythema migrans, caused by a cutaneous *B. burgdorferi* infection [54, 55]. Approximately 5% of untreated patients will develop carditis (e.g., heart block), about 10% will develop neurologic manifestations such as meningitis, cranial nerve palsy or radiculopathy, about 60% will develop arthritis [54], and about 20% of patients do not produce any subsequent clinical manifestations. The variability in clinical indicators among patients could result from individual differences or differences among the strains of *B. burgdorferi* that initiate the infections. Strains of *B. burgdorferi* can be classified into subtypes based on various typing methods. Increasing evidence suggests that certain subtypes are more likely to cause hematogenous dissemination than others [56]. Those facts that BmpA can be expressed from three independent transcripts *bmpA, bmpAbmpB* and *bmpCbmpA*, and that leader peptide is located in front of *bmpA* transcript and can regulate *bmpA* translation, suggest that differences in BmpA expression can be involved in virulence strain diversity.

## Acknowledgment

This research was done as independent research project.

## Constructs

a) *bmpA*_*L*_*::gfp*
b) *bmpA*_*L*_(ΔSD_1_)::*gfp*
c) *bmpA*_*L*_(ΔSD_2_)::*gfp*
d) *bmpA*_*L*_(33bp*bmpA*)::*gfp*
e) *bmpA*_*L*_(ΔSD_1_)33bp*bmpA*::*gfp*
f) *bmpA*_*L*_ SD_1_::*gfp*
g) *bmpA*_*L* stop_::*gfp*
h) *bmpA*_*L*ochre17_::*gfp*
i) *bmpA*_*L* mutated conserve_33bp*bmpA*::*gfp*
j) *bmpA*_*L* mutated variable_33bp*bmpA*::*gfp*

